# Environmental harshness is associated with lower investment in collective actions

**DOI:** 10.1101/663518

**Authors:** N. Lettinga, P.O. Jacquet, J-B. André, N. Baumard, C. Chevallier

## Abstract

Although humans cooperate universally, there is variability across individuals, times and cultures in the amount of resources people invest in cooperative activities. The origins of such variability are not known but recent work highlights that variations in environmental harshness may play a key role. A growing body of experimental work in evolutionary psychology suggests that humans adapt to their specific environment by calibrating their life-history strategy. In this paper, we apply structural equation models to test the association between current and childhood environmental harshness, life-history strategy and adult cooperation in two large-scale datasets (the World Values Survey and the European Values Study). The present study replicates existing research linking a harsher environment (both in adulthood and in childhood) with a modulated reproduction-maintenance trade-off and extends these findings to the domain of collective actions. Specifically, we find that a harsher environment (both in adulthood and in childhood) is associated with decreased involvement in collective action and that this association is mediated by individuals’ life-history strategy.

## 1. Introduction

Evolution by natural selection gave organisms the ability to flexibly adapt their behaviors to different environments. Human and non-human animals use this behavioral flexibility in order to maximize their reproductive success (Krebs & Davies, 2009). Life-history theory is the branch of evolutionary theory that deals with the way in which organisms allocate energy to different functions (i.e., growth, body maintenance, mating, reproduction) and with the impact of the local environment on the optimal allocation balance (Schaffer, 1983). In the past decades, a growing body of experimental work demonstrated that organisms calibrate their behavioral strategies to the specific circumstances in which they live. In humans, higher local mortality appears to affect the way in which individuals deal with the reproduction-maintenance trade-off (Ellis et al., 2009; Jasienska et al., 2017; Nettle, 2010; Promislow & Harvey, 1990), such that, in industrial and post-industrial societies, harsher conditions lead to faster growth, earlier reproduction, increased number of offspring, and diminished investment in somatic maintenance. By contrast, more favorable circumstances are associated with longer growth, delayed reproduction, fewer offspring, and increased health efforts (Del Giudice et al., 2015; Ellis et al., 2009; Jasienska et al., 2017; Promislow & Harvey, 1990; Reznick & Endler, 1982).

The evolutionary rationale behind these phenomena is that investing in early reproduction is likely adaptive when there is a high risk of dying young or being incapacitated early in life (Belsky et al., 2012, 1991, 2010; Frankenhuis et al., 2013). In humans, an important cue of experienced harshness is the degree of resource scarcity characterizing individuals’ households and neighborhoods. Indeed, greater resource scarcity correlates with virtually all forms of morbidity and mortality, indicating that people who experienced scarcity also experienced greater exposure to disease, disability and death (Adler et al., 1993; Chen et al., 2002; Chetty et al., 2016). In line with this idea, correlational research in humans has shown that early childhood environmental harshness is associated with decreased investment in health and an accelerated reproductive schedule (Brumbach et al., 2009; Mell et al., 2018; Nettle, 2011). Causal experiments in non-human animals have also demonstrated that experiencing harshness during the juvenile period has an impact on somatic and reproductive strategies (Reznick & Endler, 1982; Veenema, 2009) and natural experiments in humans have confirmed the causal influence of growing conditions on adult outcomes (Akee et al., 2018).

More recently, researchers have broadened their focus to investigate whether individuals’ ecology also has an impact on psychological traits. The most recent review on the topic highlights that harsh conditions are indeed associated with a behavioral constellation of behaviors and psychological traits that are contextually appropriate (Pepper & Nettle, 2017). Specifically, harsher ecologies tend to be associated with psychological traits such as an immediate reward orientation and a shorter time horizon (Bulley & Pepper, 2017; Griskevicius et al., 2011). One possible reason for this is that people living in harsher and unpredictable environments are less likely to reap the benefits of deferred rewards. Specifically, people with a lower socioeconomic status (SES) tend to be less future-oriented, more pessimistic about their future and more impulsive than those with a higher SES (Adams & White, 2009; de Wit et al., 2007; Robb et al., 2009). Simpson et al. (2012) also found that individuals who had been exposed to unpredictable and rapidly changing environments early in life tended to have more sexual partners and were more risk-seeking at age 23. In addition, recent works have shown an association between current and childhood environmental harshness and trust (Hörl et al., 2016; Petersen & Aarøe, 2015), electoral behavior (Safra et al., 2017) and conformist behaviors (Jacquet et al., 2019, 2018).

We extend this body of work by looking at cooperation, which is by definition, a longterm strategy: in the short term, it is always more advantageous to reap immediate benefits by being selfish and exploitative but in the long term, it is more advantageous to invest in cooperation so as to reap longer-term direct and indirect benefits (such as increased social reputation) (Axelrod & Hamilton, 1981; Baumard et al., 2013; Sjåstad, 2017; Trivers, 1971). Although humans have a moral instinct and cooperate universally, there is variability across individuals, times and cultures in the amount of resources people invest in cooperative activities (Alesina & La Ferrara, 2002; Banfield, 1967; Fukuyama, 1995; Hechter, 1988; Inglehart & Welzel, 2005). The origins of such variability remain unknown but recent work highlights that variations in environmental harshness may play a key role. In an interesting study by Nettle et al. (2011) for instance, cooperative behavior was compared across two neighborhoods with different levels of SES but similar other characteristics (e.g., total population, median age, immigration levels). They found that people living in lower SES neighborhoods were less generous in economic games, had lower social capital and showed more antisocial behavior. Safra et al. (2016) replicated this finding in 6 to 7 year-old children living in neighborhoods differing on overall SES but not other characteristics. They found that children living in harsher environments behaved less prosocially towards strangers in the context of a dictator game adapted to children. Similarly, Korndörfer et al. (2015) and Schmukle et al. (2019), using large and representative international samples, also found predominantly positive correlations between social class and prosociality (e.g., more charitable, more trusting and more helpful). McCullough et al. (2013) found that childhood environmental harshness predicts cooperative behavior in adult life. Specifically, childhood exposure to harsh environments in boys (e.g., family neglect, conflict, violence, crime) was positively correlated to exploitation of a cooperative partner and retaliation to a defective partner in an iterated prisoner’s dilemma. However, in two correlation studies involving around 500 participants each, Wu et al. (2017) found that early life environment (i.e., childhood SES and childhood unpredictability) correlated with life-history strategy (as assessed with the Mini-K and High-K strategy scale) but not with cooperation in economic games.

The aforementioned studies have several limitations. First, sample size in these existing studies is often relatively small, and not representative of the general population. Second, the High-K Strategy Scale has recently been criticized by Copping et al. (2014) for lack of construct validity. Furthermore, Olderbak et al. (2014) found that the High-K Strategy Scale and the Mini-K scale did not significantly correlate with measures of mating effort (an important life-history trait). If these scales do not correlate with such an important indicator of life-history strategy, it raises questions about the validity of these scales. Third, cooperation in economic games has recently been criticized for bearing no relationship with actual behavior (Galizzi & Navarro-Martinez, 2019). Extending on these previous studies, the present study aims to overcome these limitations. First, large-scale datasets with representative samples of respondents are used. Second, we exploit indicators of life-history strategies which come as close as possible to testing the reproduction-maintenance trade-off (i.e., respondents’ health status, respondents’ number of children, respondents’ age at their first child’s birth, Ellis et al., 2009). Third, variables reflecting investment in actual collective actions – i.e. behaviors that involve concrete costs for the agent and have benefits for others – were used as proxies for cooperation rather than self-reported attitudes.

In this paper, we thus test the relationship between current and childhood environmental harshness and people’s propensity to invest in collective actions (e.g., volunteering, activism, etc.). We further test the hypothesis that variability in cooperative behavior is mediated by differences in life-history strategy. Our specific hypotheses were that (1) current environmental harshness is associated with lower investment in collective actions, and this effect is mediated by variations in individuals’ reproduction-maintenance trade-off; (2) childhood environmental harshness is associated with lower investment in collective actions, and this effect is mediated by variations in individuals’ reproduction-maintenance trade-off. These hypotheses were tested by analyzing two large datasets that provide representative samples of respondents: the World Values Survey and the European Values Study. We used the World Values Survey to test hypothesis 1 concerning current environmental harshness. The advantage of this dataset is that it includes non-European countries and in particular poorer countries. We used the European Values Study to test hypothesis 2 concerning childhood environmental harshness. Using two independent datasets allowed us to externally replicate our results and to test the robustness of the association between harshness and cooperation. In addition, we simulated internal replication by randomly dividing each individual dataset into two subsamples of similar size: a discovery sample and a replication sample. The discovery sample was used to design and test the model. We then pre-registered the analysis plan in the Open Science Framework. The replication sample was used to check the robustness of the model.

## 2. Study 1: Current harshness and collective action in the World Values Survey

We tested the association between current environmental harshness, life-history strategy and collective action by applying structural equation modeling on the World Values Survey dataset (World Values Survey Longitudinal Data File 1981-2014 (World 1981-2014), 2015). The World Values Survey is an independent large-scale sociological survey exploring people’s values and beliefs across countries and times. The World Values Survey has relied on social scientists since 1981 to collect data from large representative samples of the populations from almost 100 countries. In this study, we used the fourth wave of the survey for which data was collected between 1999 and 2004. We focused on the fourth wave because it includes the same collective action variables as the ones included in the European Values Study, which we used in the second study reported in this paper.

### 2.1 Methods

#### 2.1.1 Respondents

59.030 respondents living in 40 different countries are included in wave 4 of the World Values Survey. However, some respondents could not be included in our analyses. First, one of the variables used to model life-history strategy is only relevant for respondents with children, which led us to focus on the 41.799 respondents who report having children. Second, we removed countries for which the cooperation variables had not been filled out, which reduced the dataset to 25.670 respondents living in 27 different countries. We randomly split this sample in a discovery subsample (12.962 respondents) and a replication subsample (12.708 respondents). In both subsamples, we excluded participants with too many empty cells (more than two standard deviations from their respective subsample mean) which led to 12.083 respondents for the discovery sample and 11.888 respondents for the replication sample. The final discovery sample after listwise deletions included 10.829 respondents (females: N = 5.633; mean age = 45 ± 15 sd; mean number of respondents per country = 417 ± 150 sd) and the final replication sample after listwise deletions included 10.617 respondents (females: N = 5.589; mean age = 45 ± 14 sd; mean number of respondents per country = 408 ± 156 sd).

#### 2.1.2 Variables

We selected variables tapping three constructs: current environmental harshness, life-history strategy and collective action. Current environmental harshness was modeled using the respondent’s current income level (scale 1-10: the higher the score the lower the income level) (Adler et al., 1993; Chen et al., 2002; Chetty et al., 2016; Griskevicius et al., 2011).

Life-history strategy was modeled by constructing a latent variable based on “subjective health status” (scale 1-5: the higher the score the better the respondents’ reported health) and “number of children”. These items were chosen because they approximate the reproduction-maintenance trade-off (Ellis et al., 2009).

Cooperation was modeled by constructing a latent variable. In articles using the World Values Survey and the European Values Study (Atkinson & Bourrat, 2011; Weeden & Kurzban, 2013), cooperation is often assessed using questions tapping cooperative morality (e.g., the respondent’s opinion about claiming government benefits to which you are not entitled; avoiding a fare on public transport; cheating on taxes if you have a chance). Such questions do no test actual cooperation, but rather people’s moral judgments. Importantly, moral judgments are not systematically aligned with actual behavior. For instance, individuals who claim that it is wrong to cheat on taxes may actually cheat themselves. Individuals living in low trust societies are indeed more likely to cheat because they think that others are cheating (Schroeder et al., 2014) and they are also more likely to be in favor of harsh punishments for cheaters because they do not trust that cheating can be avoided by means other than harsh punishments (McCullough et al., 2013).

To address this problem, we chose to focus on respondents’ actual investment in collective actions. We scanned the World Values Survey to identify variables that had previously been found to successfully predict real-life cooperation (Korndörfer et al., 2015) and that tapped actual behaviors involving concrete costs for the agent and benefits for others. Two sets of variables met our a priori defined criteria: how much someone volunteers (scale 0-14) and how politically active someone is (scale 0-4) (items are detailed in table 1 of the Supplementary Materials). Although these measures are limited in their own way (for instance, many cooperative interactions happen outside the activities described in these items), these items have the advantage of being the most concrete instances of true investment in collective goods available in the World Values Survey. Volunteering was calculated by taking the sum of 14 separate items that indicate if respondents volunteer for different groups (e.g., religious organizations, trade unions and sport groups). Political action was calculated by taking the sum of four separate items that indicate if respondents take part in different political actions (e.g., signing a petition, joining boycotts and attending lawful demonstrations).

**Table 1:**
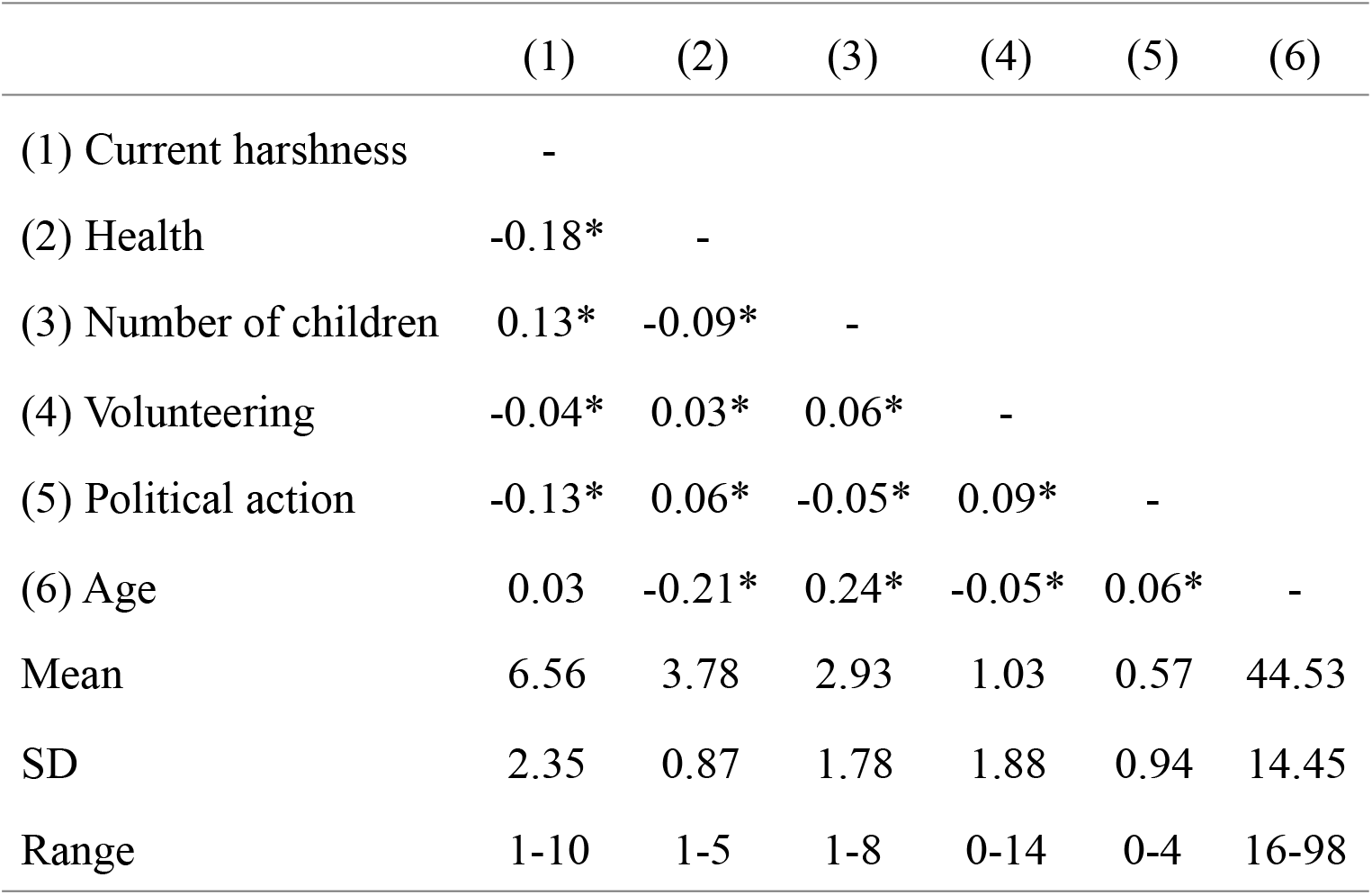
World Values Survey descriptive statistics and correlations. * *p* < 0.05.

#### 2.1.3 Model specification and fit

First, life-history strategy was regressed on current environmental harshness. Second, collective action was regressed on life-history strategy and current environmental harshness. The latent variable “life-history strategy” and the latent variable “collective action” were scaled by fixing variance to 1. This resulted in the structural equation model that is presented in figure 1. Given that cooperation is affected by age, we included age as an auxiliary variable to control for its effect on the collective action variables. Freund & Blanchard-Fields (2014), for example, found that older adults report valuing contributions to the public good more positively and are more likely to behave altruistically than younger adults. Furthermore, age is also used as an auxiliary variable to control for its effect on the “subjective health status” variable and on the “number of children” variable. The reason for this is that, all else being equal, the older you get the poorer your health is likely to be and the more children you are likely to have.

**Figure 1:**
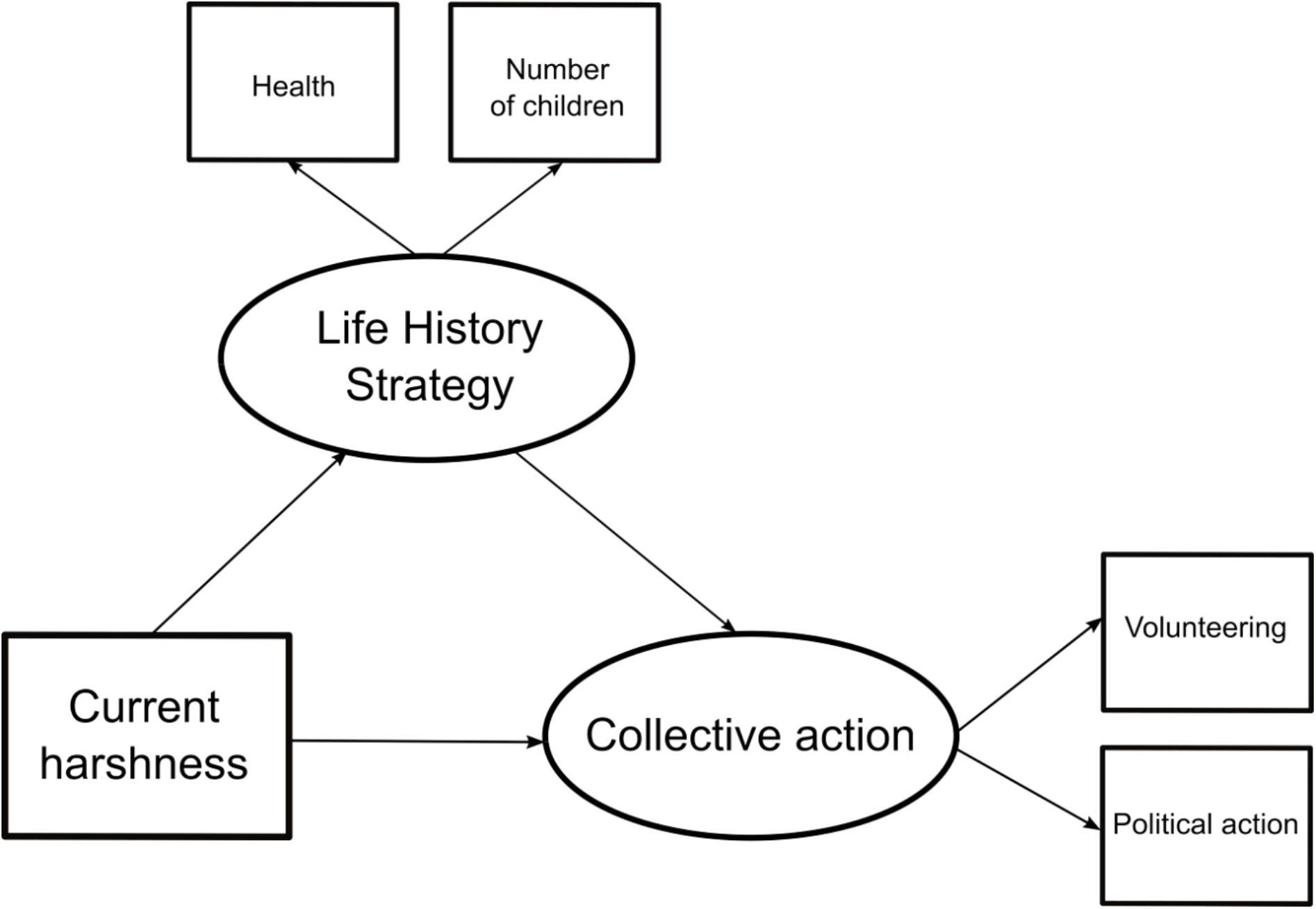
World Values Survey hypothesized structural equation model

Hooper et al. (2008) recommend determining model fit by using X^2^, Comparative Fit Index (CFI), Root Mean Square Error of Approximation (RMSEA), and Standardized Root Mean Square Residual (SRMR). A good fit is assumed if the indices are close to the following values: X^2^ p-value > 0.05, CFI > 0.95, RMSEA p-value < 0.07 and SRMR < 0.08. It is important to note however, that the large sample size of our study prevents us from interpreting X^2^ as evidence for a discrepancy between the sample and the model-implied covariance matrix. The chi-square statistic is indeed known to be particularly sensitive to sample size, which can lead models fitted on large samples to be systematically rejected (Schermelleh-Engel et al., 2003). Furthermore, the X^2^ assumes normality (McIntosh, 2007) and the deviation from normality characterizing some of our variables might result in the rejection of the model even when the model is properly specified. Therefore, only the scaled versions of the CFI, RMSEA and SRMR are reported, which eliminates the issues of sample size dependency and non-normality of continuous and categorical data (Kline, 2015).

#### 2.1.4 Analyses

We first carried out our analyses on the discovery sample; we then pre-registered our analysis plan on the Open Science Framework (the link is not shown because of the double-blind review process); finally, we applied the pre-registered analyses on the replication sample.

All statistical analyses were carried out in R 3.4.4 (https://www.r-project.org/) with R Studio 1.1.456. The SEM model was fitted using the R packages *lavaan* (Rosseel, 2012). The WLSMV estimator was used for its robustness to departures from normality (Rosseel, 2012).

When data was missing, we conducted listwise deletions. Although our variables of interest showed overall low percentages of missing responses (ranging from 0 to 13%), an additional analysis was done on the replication sample with imputed data. The results of this analysis are consistent with the results obtained using listwise deletions (see Supplementary Materials).

For the mediation analyses the bootstrap method developed by Preacher & Hayes (2008) was used, which is recommended by MacKinnon et al. (2004). This is a non-parametric resampling test. The main feature of this test is that it does not rely on the assumption of normality. Bootstrapping estimates the upper limit and the lower limit of the confidence intervals of an indirect effect. We computed bootstrapped 95% confidence intervals (1000 bootstrap samples).

### 2.2 Results

#### 2.2.1 Correlation matrix and descriptive statistics

Descriptive statistics and the correlation matrix for the variables included in the SEM can be found in table 1.

#### 2.2.2 Model fit

The scaled CFI value (0.960), the scaled RMSEA value (0.047) and the scaled SRMR value (0.018) are consistent with a close-fitting model. Therefore, the approximate fit indices reveal no strong misspecification for this model.

#### 2.2.3 Measurement model

The standardized regression weights can be found in figure 2 (the full results can be found in table 2 of the Supplementary Materials). “Subjective health status” (UnStd c = −0.15 (0.03), *p* < 0.001, Std c = −0.24) and “number of children” (UnStd c = 0.22 (0.05), *p* < 0.001, Std c = 0.17) loaded significantly on the life-history latent variable. The pattern of covariation follows our predictions: greater values on the life-history strategy latent construct indicate a strategy involving poorer reported health and higher number of children. Hence, the latent life-history construct is consistent with prior studies (Brumbach et al., 2009; Mell et al., 2018).

**Figure 2:**
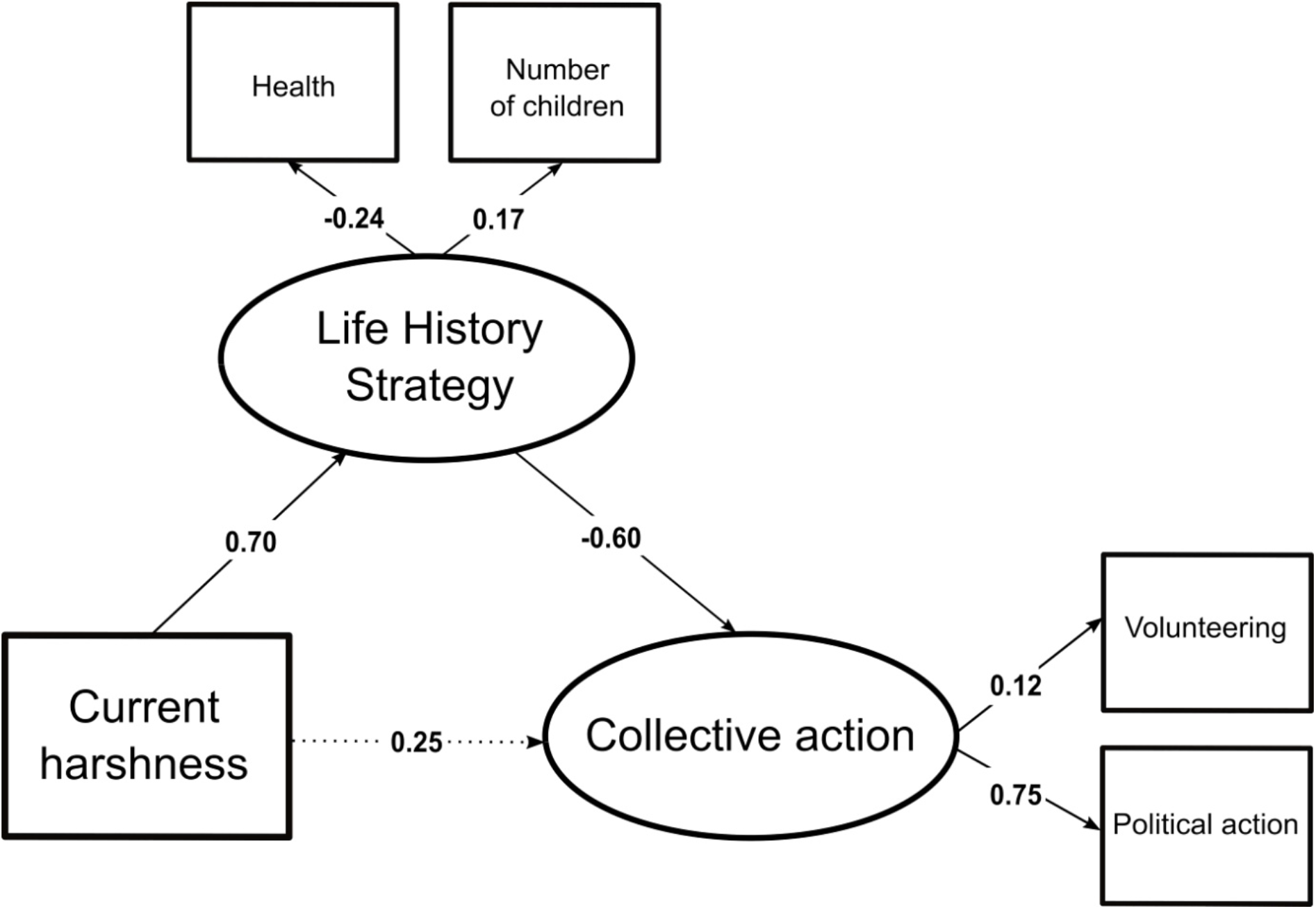
World Values Survey standardized parameter values estimated by the structural equation model. Significant paths at the 5% level are represented with a bold arrow.

**Table 2:** European Values Study descriptive statistics and correlations. * *p* < 0.05.

|  | (1) | (2) | (3) | (4) | (5) | (6) | (7) | (8) | (9) | (10) | (11) |
| --- | --- | --- | --- | --- | --- | --- | --- | --- | --- | --- | --- |
| (1) Parental education | - |  |  |  |  |  |  |  |  |  |  |
| (2) Parents problem making end meet | 0.28* | - |  |  |  |  |  |  |  |  |  |
| (3) Parents problem replacing things | 0.24* | 0.71* | - |  |  |  |  |  |  |  |  |
| (4) Death of father | 0.07* | 0.11* | 0.10* | - |  |  |  |  |  |  |  |
| (5) Death of mother | 0.02* | 0.01 | 0.01 | 0.03* | - |  |  |  |  |  |  |
| (6) Health | -0.16* | -0.16* | -0.18* | -0.06* | -0.02* | - |  |  |  |  |  |
| (7) Age 1st birth | -0.06* | -0.08* | -0.08* | -0.02 | -0.01 | 0.11* | - |  |  |  |  |
| (8) Number of children | 0.19* | 0.10* | 0.07* | 0.02 | 0.03* | -0.08* | -0.18* | - |  |  |  |
| (9) Volunteering | -0.03* | -0.02* | -0.03* | -0.01 | 0.00 | 0.1* | 0.05* | 0.04* | - |  |  |
| (10) Political action | -0.1* | -0.07* | -0.11* | -0.02* | 0.00 | 0.15* | 0.13* | -0.01 | 0.12* | - |  |
| (11) Age | 0.31* | 0.16* | 0.13* | 0.11* | 0.05* | -0.33* | 0.1* | 0.24* | 0.00 | -0.01 | - |
| Mean | 5.49 | 2.40 | 2.34 | 0.05 | 0.02 | 3.56 | 25.18 | 2.21 | 0.38 | 0.61 | 51.96 |
| SD | 1.99 | 1.17 | 1.17 | 0.22 | 0.13 | 0.96 | 5.18 | 1.12 | 1.21 | 0.96 | 15.00 |
| Range | 1-8 | 1-4 | 1-4 | 0-1 | 0-1 | 1-5 | 12-83 | 1-13 | 0-15 | 0-5 | 18-108 |

“Volunteering” (UnStd c = 0.20 (0.04), *p* < 0.001, Std c = 0.12) and “political action” (UnStd c = 0.63 (0.16), *p* < 0.001, Std c = 0.75) loaded significantly on the collective action latent variable, whose greater values indicate higher investments in political activities and, to a lesser extent, higher investments in volunteering activities.

#### 2.2.4 Structural model

Figure 2 shows that a harsher current environment (for which current income is a proxy) is associated with differences in life-history strategies (UnStd c = 0.42 (0.10), *p* < 0.001, Std c = 0.70), which is itself associated with lower involvement in collective action (UnStd c = −0.48 (0.19), *p* < 0.05, Std c = −0.60). In addition, the direct effect between current environmental harshness and adult involvement in collective action is not significant (UnStd c = 0.12 (0.11), *p* = 0.26, Std c = 0.25). In line with our first hypothesis, the effect of current environmental harshness on adult involvement in collective action is mediated by life-history strategy (indirect effect: UnStd c = −0.003, bootstrapped ci lower = −0.005, bootstrapped ci upper = −0.001, *p* < 0.001).

#### 2.2.5 Conclusions

Our prediction was that people who currently live in harsher environments invest less in collective action and that this effect is mediated by modulations of the reproduction-maintenance trade-off. Data from the World Values Survey confirm this hypothesis and show that the association between current environmental harshness and adult involvement in collective action is mediated by life-history strategies. This result is compatible with at least two (not mutually exclusive) interpretations. One interpretation is that lower involvement in collective action is the result of a flexible behavioral adjustment in response to short-term variations in participants’ resource level. Indeed, humans routinely adjust their investments in collective action in response to short-term changes in their environment: they cooperate less if they feel stressed (Riis-Vestergaard et al., 2018) or more if they feel observed (Charness & Gneezy, 2008; Franzen & Pointner, 2012). Another interpretation is that cooperation variations are in part guided by a longer-term calibration that affects a constellation of behaviors, including cooperation. Although testing the impact of long-term influences requires a properly causal design, a first step is to look at associations between childhood environmental harshness and adult levels of cooperation. Our prediction was that adult cooperation would be associated with childhood environmental harshness.

## 3. Study 2: Childhood harshness and collective action in the European Values Study

We tested the association between childhood environmental harshness, life-history strategy and adult investment in collective actions by applying structural equation modeling on the European Values Study dataset (European Values Study Longitudinal Data File 1981-2008 (EVS 1981-2008), 2015). The European Values Study is an independent large-scale sociological survey exploring people’s values and beliefs across countries and times. The European Values Study has relied on social scientists since 1981 to collect data from large representative samples of the populations from 47 European countries. In this study, we used the fourth wave of the survey for which data was collected between 2008 and 2010. We focused on the fourth wave because it includes the same collective action variables as the World Values Survey, which gives us the opportunity to replicate our initial findings on an independent sample. Furthermore, unlike wave 4 in the World Values Survey, wave 4 in the European Values Study provides access to variables assessing the quality of the environment experienced by respondents during their childhood. Specifically, it includes variables indexing the socio-economic status of the respondents’ parents (i.e., resource scarcity of the family household, education of the parents), but also variables indexing respondents’ direct exposure to extrinsic mortality during childhood (death of the father, death of the mother). Finally, it allows for the inclusion of an additional and highly representative parameter of life-history strategy: respondents’ age at their first child’s birth. Age at first birth is an interesting index of the respondents’ investment in reproduction, and a key aspect of an individual’s life-history strategy. For instance, it has been shown that women’s age at first birth is younger when extrinsic mortality risks are higher (Low et al., 2008). One possible explanation for this acceleration of reproductive scheduling is that it offsets the fitness cost of high extrinsic mortality by increasing the chance of reproducing before death (Bulley & Pepper, 2017).

### 3.1 Methods

#### 3.1.1 Respondents

66.281 respondents living in 46 different countries are included in wave 4 of the European Values Study. However, some respondents could not be included in our analyses. Two of the variables used to model life-history strategy were only relevant for respondents with children, which led us to focus on the 45.624 respondents who report having children. We randomly split this sample in a discovery subsample (22.570 respondents) and a replication subsample (23.054 respondents). In both subsamples, we excluded participants with too many empty cells (more than two standard deviations from their respective subsample mean) which led to 20.755 respondents for the discovery sample and 21.156 respondents for the replication sample. The final discovery sample after listwise deletions included 16.790 respondents (females: N = 9.837; mean age = 52 ± 15 sd; mean number of respondents per country = 382 ± 126 sd) and the final replication sample after listwise deletions included 17.087 respondents (females: N = 9.957; mean age = 52 ± 15 sd; mean number of respondents per country = 371 ± 146 sd).

#### 3.1.2 Variables

We selected variables tapping three constructs: childhood environmental harshness, life-history strategy and collective action. Childhood environmental harshness was modeled by calculating a composite score including the following items: “death of the father before age 16” (scale 0-1), “death of the mother before age 16” (scale 0-1), “parental education” (scale 1-8: the higher the score the lower the educational level of the parents), “parents had problems replacing things” (scale 1-4: the higher the score the more difficulty replacing things) and “parent’s had problems making end meet” (scale 1-4: the higher the score the more difficulty making ends meet). Based on the recommendations of Brumbach et al. (2009), childhood environmental harshness was modeled as an emergent variable rather than a reflective latent variable. This was done because harsher childhood events can be seen as risk factors that are not necessarily correlated with one another. However, they all contribute to the cumulative probability of developing a particular outcome (in our case a particular reproductive or somatic maintenance strategy). For example, having been exposed to the death of a parent, lower parental education or lower parental SES are three events that increase the probability of altering individuals’ life-history strategy, but that might occur independently.

As in study 1, life-history strategy was modeled by constructing a latent variable based on “subjective health status” (scale 1-5: the higher the score the better the respondents’ reported health) and “number of children”, to which the new indicator “age at first birth” was added. “Age at first birth” was calculated from the birth year of the respondent and the reported birth year of their first child.

The latent construct “collective action” was estimated from the very same indicators as in study 1. The only difference is that scores in “volunteering” and “political action” were calculated on the basis of 15 and 5 items respectively (instead of 14 and 4 items in study 1) (items are detailed in table 3 of the Supplementary Materials).

#### 3.1.3 Model specification and fit

The model’s structure was the same as the one specified in study 1, with two notable differences. First, current environmental harshness was replaced by childhood environmental harshness, modeled as a composite variable. Composite variables need to be scaled for identification purposes by fixing the coefficient of one of the causal indicators (Mell et al., 2018). Childhood environmental harshness was scaled by setting the path from “parental education” to 1. Second, the correlation between the residual errors of “age at first birth” and “number of children” is included in the model. The reason for this is that, all else being equal, the sooner you have children, the more children you can have. This correlation is not captured by the latent variable life-history strategy and is therefore included separately; these two variables indeed covary mechanically (Mell et al., 2018). This resulted in the structural equation model that is presented in figure 3. Model fit was determined in the same manner as in study 1.

**Figure 3:**
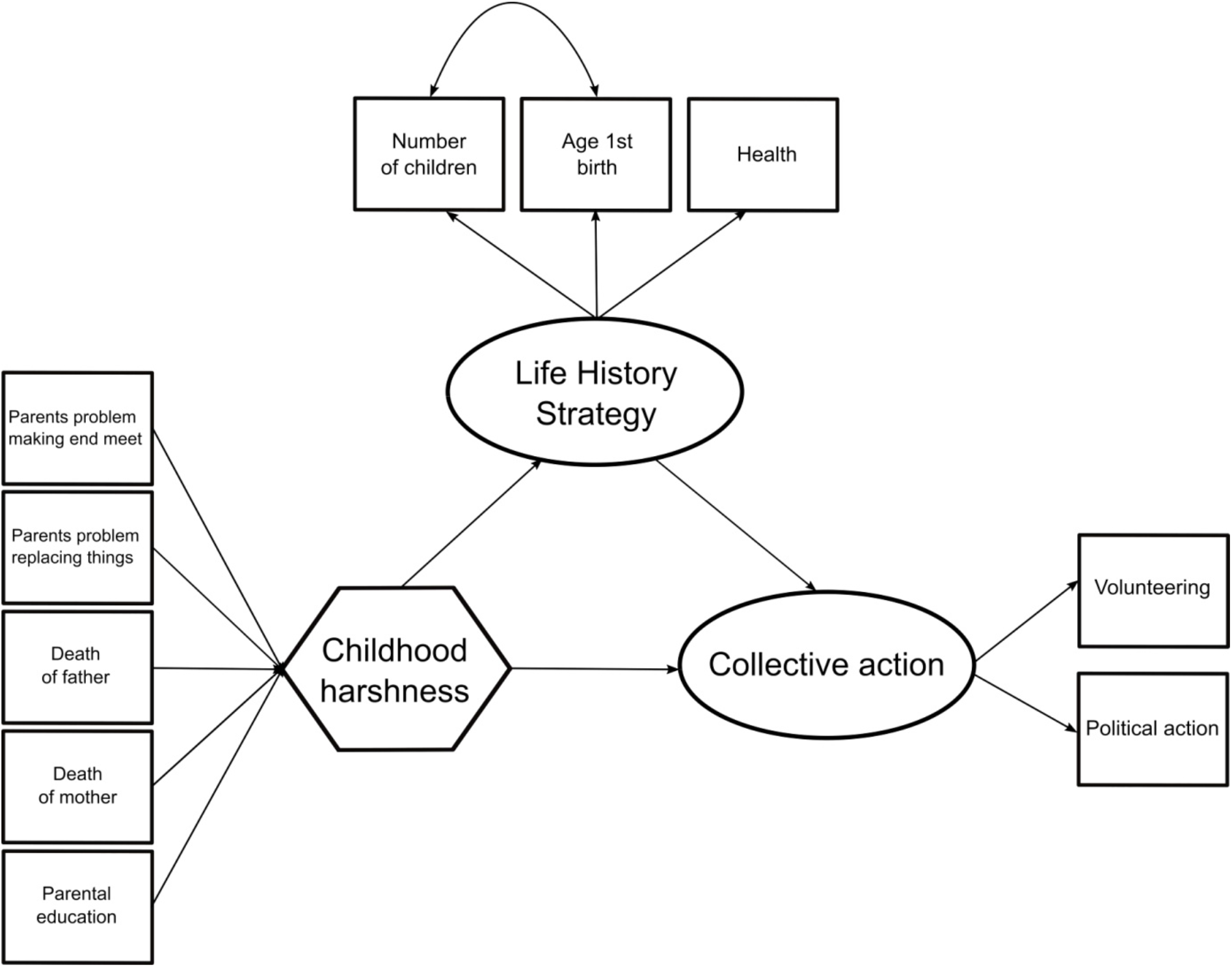
European Values Study hypothesized structural equation model

#### 3.1.4 Analyses

We first carried out our analyses on the discovery sample; we then pre-registered our analysis plan on the Open Science Framework (the link is not shown because of the double-blind review process); finally, we applied the pre-registered analyses on the replication sample.

All statistical analyses and the method for mediation analyses were similar to those reported in study 1. When data was missing, we conducted listwise deletions (as in study 1). Although our variables of interest showed overall low percentages of missing responses (ranging from 0 to 13%), an additional analysis was done on the replication sample with imputed data. The results of this analysis are consistent with the results obtained using listwise deletions (see Supplementary Materials).

### 3.2 Results

#### 3.2.1 Correlation matrix and descriptive statistics

Descriptive statistics and the correlation matrix for the variables included in the SEM can be found in table 2.

#### 3.2.2 Model fit

The scaled CFI value (0.900), the scaled RMSEA value (0.040) and the scaled SRMR value (0.018) are consistent with a close-fitting model. Therefore, the approximate fit indices reveal no strong misspecification for this model.

#### 3.2.3 Measurement model

The standardized regression weights can be found in figure 4 (the full results can be found in table 4 of the Supplementary Materials). Childhood environmental harshness as a composite is driven by two main contributors: “parents had problems replacing things” (UnStd c = 2.37 (0.26), *p* < 0.001, Std c = 0.69) and “death of the father before age 16” (UnStd c = 1.65 (0.70), *p* < 0.05, Std c = 0.09). “Death of the mother before age 16” and “parents had problems makings ends meet” did not load significantly on childhood environmental harshness.

**Figure 4:**
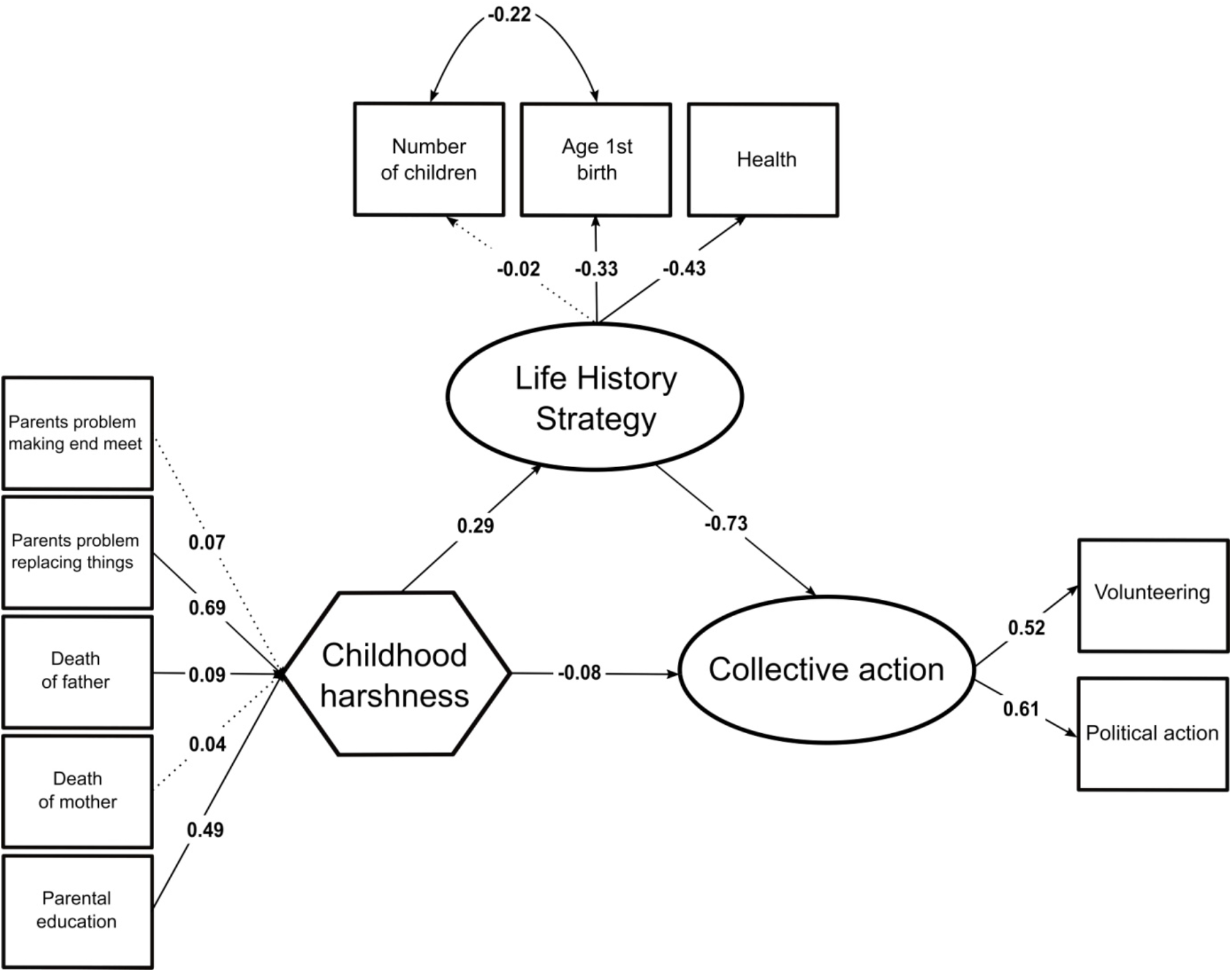
European Values Study standardized parameter values estimated by the structural equation model. Significant paths at the 5% level are represented with a bold arrow.

“Subjective health status” (UnStd c = −0.39 (0.02), *p* < 0.001, Std c = −0.43) and “age at first birth” (UnStd c = −1.60 (0.07), *p* < 0.001, Std c = −0.33) loaded significantly on the life-history latent variable. As in study 1, the pattern of covariation follows our predictions: greater values of the life-history strategy latent construct indicate a strategy involving poorer reported health and a younger age at first child’s birth. “Number of children” did not load significantly on childhood environmental harshness. However, inspection of the estimated covariance shows that “number of children” is not independent but covaries with “age at first birth” in the expected way (UnStd c = −1.15 (0.04), *p* < 0.001, Std c = −0.22), an observation that is in line with existing findings (Mell et al., 2018). Hence, the latent life-history construct is fairly consistent with prior studies (Brumbach et al., 2009; Mell et al., 2018).

“Volunteering” (UnStd c = 0.34 (0.02), *p* < 0.001, Std c = 0.52) and “political action” (UnStd c = 0.40 (0.03), *p* < 0.001, Std c = 0.61) loaded significantly on the collective action latent variable, whose greater values indicate higher investments in both volunteering and political activities.

#### 3.2.4 Structural model

Figure 4 shows that a harsher childhood environment is associated with differences in life-history strategies (UnStd c = 0.08 (0.01), *p* < 0.001, Std c = 0.29), which is itself associated with lower adult involvement in collective action (UnStd c = −1.07 (0.10), *p* < 0.001, Std c = −0.73). In addition, the direct effect of childhood environmental harshness is significantly – albeit weakly – associated with adult involvement in collective action (UnStd c = −0.03 (0.01), *p* < 0.001, Std c = −0.08). This implies that a harsher childhood environment is associated with less investment in collective actions later in life, relatively independently of one’s life-history strategy. In line with our second hypothesis, a significant part of the effect of childhood environmental harshness on adult involvement in collective action was nevertheless mediated by life-history strategy (indirect effect: UnStd c = −0.003, bootstrapped ci lower = −0.003, bootstrapped ci upper = −0.002, *p* < 0.001).

#### 3.2.5 Current environmental harshness

Finally, we controlled for the effect of current environmental harshness on the life-history strategy and collective action variables. Current environmental harshness was modeled using the respondent’s current income level (scale 1-10: the higher the score the lower the income level). The inclusion of current environmental harshness leaves the model fit relatively unaffected (scaled CFI = 0.897; scaled RMSEA = 0.042; scaled SRMR = 0.017), and the magnitude of key parameters stayed significant and in the expected direction (the full results can be found in table 5 of the Supplementary Materials). Thus, when accounting for the effect of current environmental harshness, childhood environmental harshness remains a significant predictor of adult involvement in collective action via one’s life-history strategy.

#### 3.2.6 Conclusions

Our prediction was that people who grew up in harsher environments invest less in collective actions later in life and that this effect is mediated by modulations of the reproduction-maintenance trade-off. Data from the European Values Study partly confirm this hypothesis and show that the association between childhood environmental harshness and adult involvement in collective action is partly mediated by life-history strategies.

## 4. Discussion

The present study replicated existing research linking a harsher current and childhood environment with variations in individuals’ life-history strategy (Ellis et al., 2009; Mell et al., 2018). Our study extends these findings to the domain of cooperation. Specifically, we find that a harsher current and childhood environment is associated with decreased involvement in collective action and that this association is mediated by individuals’ life-history strategy.

The rationale for studying the specific effect of childhood environmental harshness is that it provides an indication that individuals’ actual cooperative strategy possibly results from long-term calibration processes. Previous works have already gathered some evidence for the existence of such early calibration in the domains of trust (Hörl et al., 2016; Petersen & Aarøe, 2015), electoral behavior (Safra et al., 2017) and conformist behaviors (Jacquet et al., 2019, 2018). Indeed, the effect of current environmental harshness on life-history strategies and cooperation might simply reflect the flexible adjustment of behavior in response to shortterm variations in local contingencies. For instance, individuals may cooperate less because they may ‘reason’ or ‘feel’ that they are currently short on money or time, and that they cannot afford to invest in their community. The present study contributes to this literature by showing that there is an association between childhood environmental harshness and adult cooperation. More generally, our findings are consistent with the idea that a harsher childhood experience leads to a constellation of behaviors adapted to harsh life conditions (Pepper & Nettle, 2017).

Even though the findings reported above are consistent with the existing literature, it is important that we highlight several limitations. First, we used people’s involvement in collective actions as a proxy for cooperation but cooperation obviously encompasses more than collective action behaviors. The reason we chose these items is that these were the only variables that met our a priori defined criteria (actual behaviors involving concrete costs for the agent and have benefits for others) in the World Values Survey and the European Values Study.

Second, our goal was to investigate if part of the variability in people’s involvement in collective action was mediated by differences in life-history strategy. Although life-history strategy is well captured in the World Values Survey model (R^2^ = 0.50), it is not well captured in the European Values Study model (R^2^ = 0.09). This may reflect the fact that life-history traits are influenced by multiple causes beyond life-history, including cultural factors like contraceptives. Such cultural factors might account for the fact that “number of children” is not significantly correlated with respondents’ life-history strategy in the European Values Study (Colleran, 2016).

More generally, the correlational nature of our data hinders the possibility of producing inferences about the causal role of harsher environments on cooperation. For example, it is possible that the relationship between environmental harshness and cooperation is reversed: harsher environments might be associated with lower cooperation because individuals who cooperate less are less likely to do well in life and more likely to end up living in a harsh environment. This alternative is plausible albeit a little harder to reconcile with the fact that we also found an association between a harsher childhood environment and adult cooperation. Another possibility is that there is an unknown variable that influences both environmental harshness and cooperation.

One way to demonstrate the causal effect of environmental harshness on cooperation would be to experimentally manipulate acute stress (Haushofer & Fehr, 2014), for example by administrating hydrocortisone (a synthetic form of cortisol). For example, Riis-Vestergaard et al. (2018) found that administering hydrocortisone increases temporal discounting over the short term. Another possibility to induce stress is to expose people to mortality cues (Griskevicius et al., 2011), e.g. in the form of news articles stating that in the local area there is an increase in violence. However, it is important to note that there are large differences between the effects of acute stress and chronic stress (Kandasamy et al., 2014). For example, acute stress has been found to increase physical arousal, learning, motivated behavior and sensation seeking. By contrast, chronic stress has been found to impair attentional control, behavioral flexibility and it can promote anxiety, depression and learned helplessness.

Another option to identify causal relationships is to analyze longitudinal data involving exogenous shocks to the individual’s environment (e.g., sudden increase of income or in contrast famine, war, etc.). There have been a handful of longitudinal studies testing the effect of stress during childhood and social behavior in adulthood using exogenous shocks of positive or negative income shocks on different behavioral outcomes. These studies have shown that higher levels of income tend to make individuals more ‘agreeable’ and more trustful (Akee et al., 2018; Hörl et al., 2016) while higher levels of violence during childhood make people more antisocial and more violent (Gangadharan et al., 2017; Miguel et al., 2011). One of the most noteworthy pieces of research in this area is a quasi-experimental study led Akee et al. (2018). In this study, American Indian children were exposed to a positive income shock following the opening of a casino that decided to allocate part of its revenue to Cherokee households. Particularly interesting is that the increase in income led to an increase in agreeableness, which correlates with cooperation and unselfishness. Hence, at least part of the effects of childhood environmental harshness on cooperation might capture long-term plastic responses to the environment.

Finally, it should be noted that our results are compatible with a range of evolutionary mechanisms. The standard mechanism put forward in the literature (and in our introduction) is that cooperation is adaptively reduced in harsher environments because individuals have a shorter life expectancy and should therefore be more short-term oriented (Nettle et al., 2011; Petersen & Aarøe, 2015). However, the idea that people focus more on the present because they have a higher probability of dying at age 60 instead of age 80 is problematic. Why should this affect their interest in cooperation today? The fact that someone is alive at age 60 or age 80 indeed has little impact on his or her probability of being alive tomorrow (Mell et al., 2017; Riis-Vestergaard & Haushofer, 2017). Another possibility is that people living in harsher environments cooperate less because they have less capital. Lower levels of resources thus make cooperation more risky because people cannot afford to lose resources and thus cannot take the risk of being cheated. Given that cooperation is an investment in social interactions, it carries benefits and costs that arise with some degree of uncertainty. Depending on individuals’ level of resources, we thus might expect different levels of investment in cooperation. Assuming marginally decreasing returns from the amounts of resources possessed, the perspective of losing even a small quantity of resources for a low-SES individual with almost no capital will indeed be dramatic compared to a rich individual with a comfortable amount of resources to buffer the losses. Hence, in addition to being less willing to cooperate overall, we can make the additional prediction that low-SES individuals should be particularly sensitive both to an increase in the probability of exploitation and to the size of potential losses compared to high-SES individuals.

To conclude this article, we would like to emphasize the relevance of this kind of research for the understanding of social institutions. Social institutions are usually analyzed through the lens of rational choice theory and are seen as the result of bargaining between self-interested social groups. For instance, when inequality is high and some groups have more power, institutions are less likely to be inclusive and democracy is less likely to appear (Acemoglu & Robinson, 2015). These kinds of theories do not take into account the fact that humans are a cooperative species, and that their willingness to cooperate varies depending on their environment. A behavioral ecology approach to cooperation by contrast, could help explain why some environments, where harshness is low, are more favorable to social institutions. These environments can indeed build on more cooperative individuals, which in turn favor inclusive institutions that are themselves associated with higher levels of economic affluence because citizens are more likely to push for social justice (Boix, 2011).

## Acknowledgements

This study was supported by the Institut d’Etudes Cognitives (ANR-10-IDEX-0001-02 FrontCog), the Institut national de la santé et de la recherche médicale (INSERM), an ERC consolidator grant (no. 8657) and an “Action Incitative” funding from the École Normale Supérieure.

