## supplementary material for "Environmental harshness is associated with lower investment in collective actions"

1. **World Values Survey collective action items**

Volunteering

Unpaid work social welfare service for elderly, handicapped or deprived people

Unpaid work religious or church

Unpaid work education, arts, music or cultural activities

Unpaid work labour unions

Unpaid work political parties or groups

Unpaid work local political action groups

Unpaid work human rights

Unpaid work environment, conservation, animal rights

Unpaid work professional associations

Unpaid work youth work

Unpaid work sports or recreation

Unpaid work women´s group

Unpaid work peace movement

Unpaid work organization concerned with health

Political action

Signing a petition

Joining in boycotts

Attending lawful/peaceful demonstrations

Joining unofficial strikes

**Supplementary table 1:** World Values Survey collective action items

2. **World Values Survey listwise deletions - full results**

| Variable | Regressed on | Unstandardized C | Standard Error | P-value | Standardized C |
| --- | --- | --- | --- | --- | --- |
| Life-history strategy | Health | -0.15 | 0.03 | 0.00 | -0.24 |
| Life-history strategy | Number of children | 0.22 | 0.05 | 0.00 | 0.17 |
| Collective action | Volunteering | 0.20 | 0.04 | 0.00 | 0.12 |
| Collective action | Political action | 0.63 | 0.16 | 0.00 | 0.75 |
| Life-history strategy | Current harshness | 0.42 | 0.10 | 0.00 | 0.70 |
| Collective action | Life-history strategy | -0.48 | 0.19 | 0.01 | -0.60 |
| Collective action | Current harshness | 0.12 | 0.11 | 0.26 | 0.25 |

**Supplementary table 2:** World Values Survey listwise deletions – full results

3. **World Values Survey imputed data results**

**Analyses**

For the imputed data twenty complete datasets were generated by fully conditional specifications for categorical and continuous data using the r package *mice* (Buuren & Groothuis-Oudshoorn, 2010). This package allows the use of different imputation methods depending on the type of variable with missing entries. Predictive mean matching was used for numeric variables, logistic regression imputation for binary data and proportional odds model for ordered categorical variables with more than two levels. The function runMI of the R package *semTools* (Contributors, 2016) was used to combine the results obtained for the 20 imputed datasets.

For the mediation analyses the bootstrap method developed by Preacher & Hayes (2008) was used, which is recommended by MacKinnon et al. (2004). This is a non-parametric resampling test. The main feature of this test is that it does not rely on the assumption of normality. Bootstrapping estimates the upper limit and the lower limit of the confidence intervals of an indirect effect. We computed bootstrapped 95% confidence intervals (1000 bootstrap samples) for each of the 20 imputed datasets and then took the average of these datasets.

**Model fit**

The scaled CFI value (0.957), the scaled RMSEA value (0.049) and the scaled SRMR value (0.018) are consistent with a close-fitting model. Therefore, the approximate fit indices reveal no strong misspecification for this model.

**Measurement model**

The standardized regression weights can be found in supplementary figure 1. “Subjective health status” (UnStd c = -0.14 (0.04), *p* < 0.001, Std c = -0.24) and “number of children” (UnStd c = 0.20 (0.05), *p* < 0.001, Std c = 0.16) loaded significantly on the life-history latent variable. The pattern of covariation follows our predictions: greater values on the life-history strategy latent construct indicate a strategy involving poorer reported health and higher number of children. Hence, the latent life-history construct is consistent with prior studies (Brumbach et al., 2009; Mell et al., 2018).

“Volunteering” (UnStd c = 0.20 (0.03), *p* < 0.001, Std c = 0.13) and “political action” (UnStd c = 0.56 (0.14), *p* < 0.001, Std c = 0.71) loaded significantly on the collective action latent variable, whose greater values indicate higher investments in political activities and, to a lesser extent, higher investments in volunteering activities.

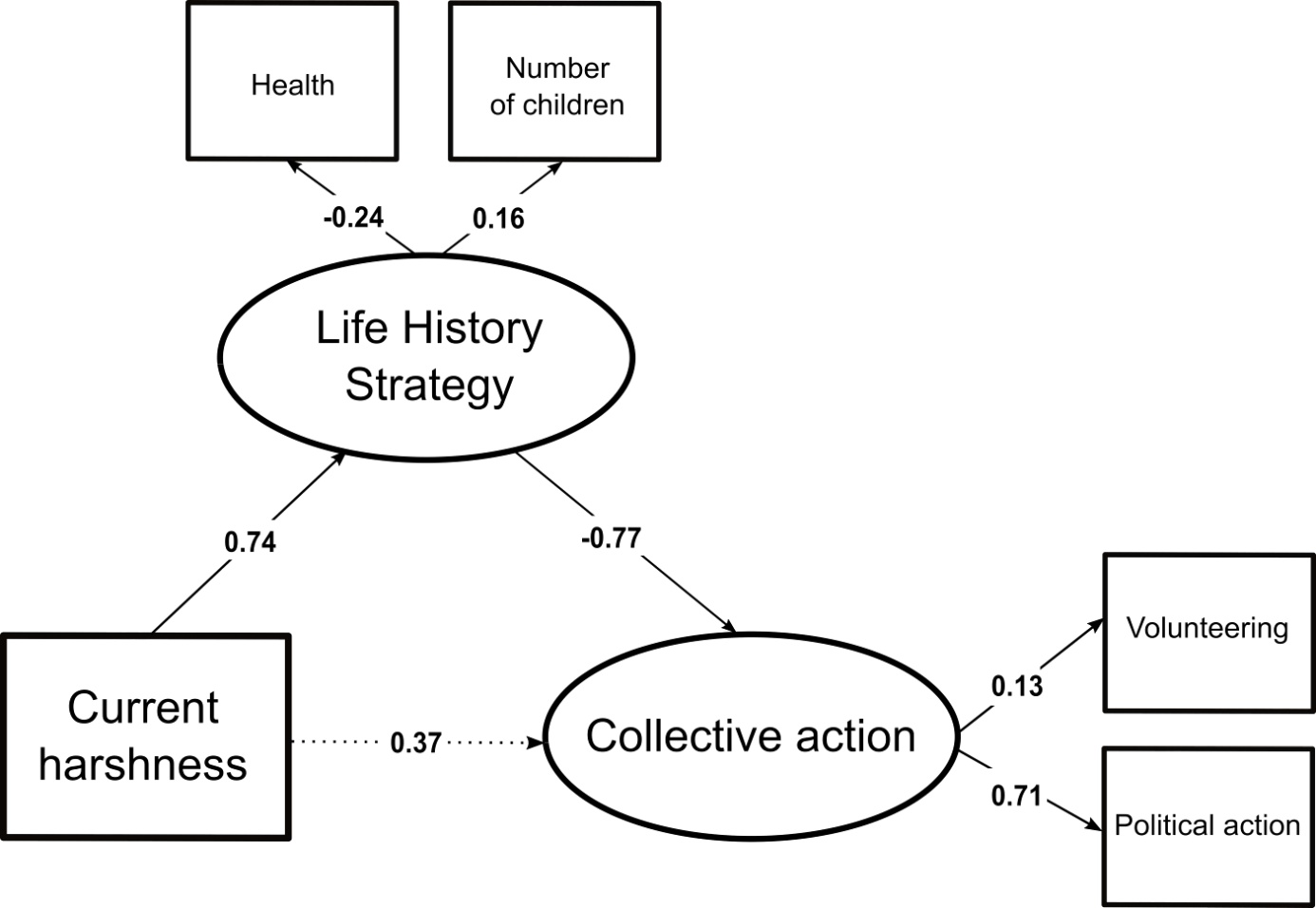

**Supplementary figure 1:** World Values Survey standardized parameter values estimated by the structural equation model. Significant paths at the 5% level are represented with a bold arrow.

**Structural model**

Supplementary figure 1 shows that a harsher current environment (for which current income is a proxy) is associated with differences in life-history strategies (UnStd c = 0.47 (0.12), *p* < 0.001, Std c = 0.74), which is itself associated with lower involvement in collective action (UnStd c = -0.62 (0.29), *p* < 0.05, Std c = -0.77). In addition, the direct effect between current environmental harshness and adult involvement in collective action is not significant (UnStd c = 0.19 (0.18), *p* = 0.30, Std c = 0.37). In line with our first hypothesis, the effect of current environmental harshness on adult involvement in collective action is mediated by life-history strategy (indirect effect: UnStd c = -0.003, bootstrapped ci lower = -0.005, bootstrapped ci upper = -0.001, *p* < 0.001).

4. **European Values Study collective action items**

Volunteering

Unpaid work social welfare service for elderly, handicapped or deprived people

Unpaid work religious or church

Unpaid work education, arts, music or cultural activities

Unpaid work labour unions

Unpaid work political parties or groups

Unpaid work local political action groups

Unpaid work human rights

Unpaid work environment, conservation, animal rights

Unpaid work professional associations

Unpaid work youth work

Unpaid work sports or recreation

Unpaid work women´s group

Unpaid work peace movement

Unpaid work organization concerned with health

Unpaid work other groups

Political action

Signing a petition

Joining in boycotts

Attending lawful/peaceful demonstrations

Joining unofficial strikes

Occupying buildings or factories

**Supplementary table 3:** European Values Study collective action items

5. **European Values Study listwise deletions - full results**

| Variable | Regressed on | Unstandardized C | Standard Error | P-value | Standardized C |
| --- | --- | --- | --- | --- | --- |
| Childhood harshness | Parental education | 1.00 | - | - | 0.49 |
| Childhood harshness | Parents problems making ends meet | 0.25 | 0.18 | 0.17 | 0.07 |
| Childhood harshness | Parents problems replacing things | 2.37 | 0.26 | 0.00 | 0.69 |
| Childhood harshness | Death of father | 1.65 | 0.70 | 0.02 | 0.09 |
| Childhood harshness | Death of mother | 1.24 | 1.06 | 0.24 | 0.04 |
| Life-history strategy | Health | -0.39 | 0.02 | 0.00 | -0.43 |
| Life-history strategy | Age at 1st birth | -1.60 | 0.07 | 0.00 | -0.33 |
| Life-history strategy | Number of children | -0.02 | 0.01 | 0.15 | -0.02 |
| Collective action | Volunteering | 0.34 | 0.02 | 0.00 | 0.52 |
| Collective action | Political action | 0.40 | 0.03 | 0.00 | 0.61 |
| Life-history strategy | Childhood harshness | 0.08 | 0.01 | 0.00 | 0.29 |
| Collective action | Life-history strategy | -1.07 | 0.10 | 0.00 | -0.73 |
| Collective action | Childhood harshness | -0.03 | 0.01 | 0.00 | -0.08 |
| Age at 1st birth | Number of children | -1.15 | 0.04 | 0.00 | -0.22 |

**Supplementary table 4:** European Values Study listwise deletions – full results

6. **European Values Study imputed data results**

**Analyses**

For the imputed data twenty complete datasets were generated by fully conditional specifications for categorical and continuous data using the r package *mice* (Buuren & Groothuis-Oudshoorn, 2010). This package allows the use of different imputation methods depending on the type of variable with missing entries. Predictive mean matching was used for numeric variables, logistic regression imputation for binary data and proportional odds model for ordered categorical variables with more than two levels. The function runMI of the R package *semTools* (Contributors, 2016) was used to combine the results obtained for the 20 imputed datasets.

For the mediation analyses the bootstrap method developed by Preacher & Hayes (2008) was used, which is recommended by MacKinnon et al. (2004). This is a non-parametric resampling test. The main feature of this test is that it does not rely on the assumption of normality. Bootstrapping estimates the upper limit and the lower limit of the confidence intervals of an indirect effect. We computed bootstrapped 95% confidence intervals (1000 bootstrap samples) for each of the 20 imputed datasets and then took the average of these datasets.

**Model fit**

The scaled CFI value (0.964), the scaled RMSEA value (0.039) and the scaled SRMR value (0.023) are consistent with a close-fitting model. Therefore, the approximate fit indices reveal no strong misspecification for this model.

**Measurement model**

The standardized regression weights can be found in supplementary figure 2. Childhood environmental harshness as a composite is driven by “parents had problems replacing things” (UnStd c = 4.41 (0.69), *p* < 0.001, Std c = 0.81), indicating that a harsher childhood environment is associated with parents having more problems replacing things. “Death of the father before age 16”, “death of the mother before age 16” and “parents had problems makings ends meet” did not load significantly on childhood environmental harshness.

“Subjective health status” (UnStd c = -0.33 (0.02), *p* < 0.001, Std c = -0.37), “age at first birth” (UnStd c = -1.17 (0.07), *p* < 0.001, Std c = -0.24) and “number of children” (UnStd c = 0.07 (0.01), *p* < 0.001, Std c = 0.07) loaded significantly on the life-history latent variable. The pattern of covariation follows our predictions: greater values of the life-history strategy latent construct indicate a strategy involving poorer reported health, a younger age at first child’s birth and higher number of children. Inspection of the estimated covariance shows that “number of children” covaries with “age at first birth” in the expected way (UnStd c = -0.94 (0.04), *p* < 0.001, Std c = -0.17), an observation that is in line with existing findings (Mell et al., 2018). Hence, the latent life-history construct is consistent with prior studies (Brumbach et al., 2009; Mell et al., 2018).

“Volunteering” (UnStd c = 0.16 (0.03), *p* < 0.001, Std c = 0.25) and “political action” (UnStd c = 0.26 (0.05), *p* < 0.001, Std c = 0.50) loaded significantly on the collective action latent variable, whose greater values indicate higher investments in both volunteering and political activities.

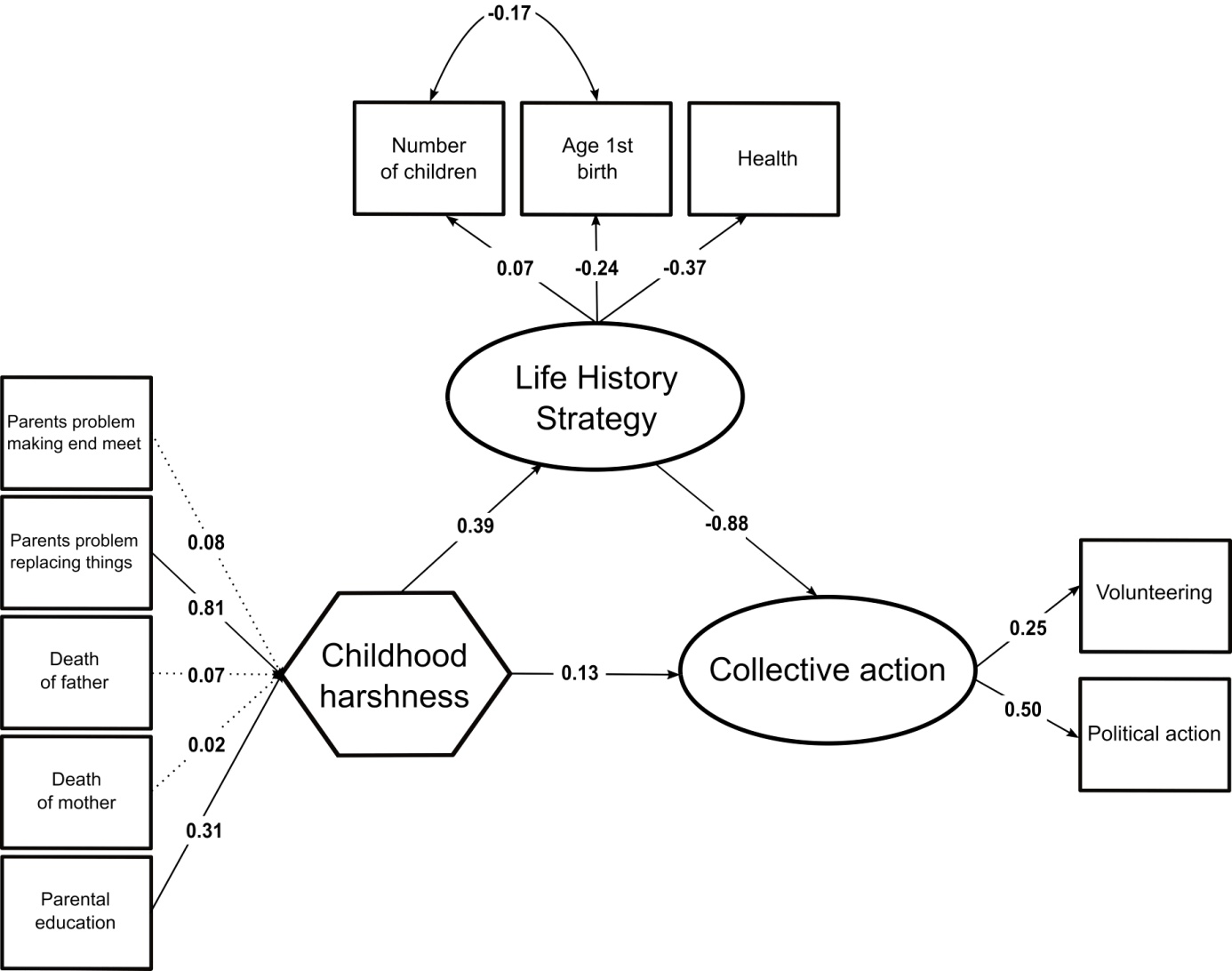

**Supplementary figure 2:** European Values Study standardized parameter values estimated by the structural equation model. Significant paths at the 5% level are represented with a bold arrow.

**Structural model**

Supplementary figure 2 shows that a harsher childhood environment is associated with differences in life-history strategies (UnStd c = 0.07 (0.01), *p* < 0.001, Std c = 0.39), which is itself associated with lower adult involvement in collective action (UnStd c = -1.50 (0.34), *p* < 0.001, Std c = -0.88). In addition, the direct effect of childhood environmental harshness is significantly – albeit weakly – associated with adult investment in collective action (UnStd c = 0.04 (0.02), *p* < 0.05, Std c = 0.13). This implies that a harsher childhood environment is associated with less investment in collective actions later in life, relatively independently of one’s life-history strategy. In line with our second hypothesis, a significant part of the effect of childhood environmental harshness on adult involvement in collective action was nevertheless mediated by life-history strategy (indirect effect: UnStd c = -0.002, bootstrapped ci lower = -0.003, bootstrapped ci upper = -0.002, *p* < 0.001).

7. **European Values Study controlled for current environmental harshness**

| Variable | Regressed on | Unstandardized C | Standard Error | P-value | Standardized C |
| --- | --- | --- | --- | --- | --- |
| Childhood harshness | Parental education | 1.00 | - | - | 0.41 |
| Childhood harshness | Parents problems making ends meet | 0.04 | 0.29 | 0.90 | 0.01 |
| Childhood harshness | Parents problems replacing things | 3.23 | 0.51 | 0.00 | 0.79 |
| Childhood harshness | Death of father | 2.15 | 1.14 | 0.06 | 0.10 |
| Childhood harshness | Death of mother | 2.16 | 1.82 | 0.24 | 0.06 |
| Life-history strategy | Health | -0.36 | 0.02 | 0.00 | -0.39 |
| Life-history strategy | Age at 1st birth | -1.44 | 0.07 | 0.00 | -0.29 |
| Life-history strategy | Number of children | -0.05 | 0.02 | 0.00 | -0.04 |
| Collective action | Volunteering | 0.35 | 0.02 | 0.00 | 0.51 |
| Collective action | Political action | 0.40 | 0.03 | 0.00 | 0.57 |
| Life-history strategy | Childhood harshness | 0.05 | 0.01 | 0.00 | 0.24 |
| Collective action | Life-history strategy | -1.04 | 0.11 | 0.00 | -0.72 |
| Collective action | Childhood harshness | -0.02 | 0.01 | 0.01 | -0.07 |
| Age at 1st birth | Number of children | -1.17 | 0.04 | 0.00 | -0.23 |

**Supplementary table 5:** European Values Study controlled for current environmental harshness
